# Zinc Boosts EGCG’s hIAPP Amyloid Inhibition Both in Solution and Membrane

**DOI:** 10.1101/401521

**Authors:** Young-Ho Lee, Yuxi Lin, Sarah J. Cox, Misaki Kinoshita, Bikash R. Sahoo, Magdalena Ivanova, Ayyalusamy Ramamoorthy

**Affiliations:** Institute for Protein research, Osaka University Yamadaoka 3-2, Suita, Osaka 565-0871, Japan.; Protein Structure Research Group, Division of Bioconvergence Analysis, Korea Basic Science Institute, Chungcheongbuk-do 28119, South Korea.; Department of Chemistry, Sookmyung Women’s University Cheongpa-ro 47-gil 100, Yongsangu, Seoul 04310, South Korea.; Biophysics and Department of Chemistry, University of Michigan Ann Arbor, MI 48109-1055, USA.; Department of Neurology, University of Michigan, Ann Arbor, MI 48109, USA

**Keywords:** epigallocatechin-gallate, islet amyloid polypeptide, type II diabetes, amyloid, protein aggregation

## Abstract

Amyloid aggregation of human islet amyloid polypeptide (hIAPP) is linked to insulin-producing islet cell death in type II diabetes. Previous studies have shown the amyloid inhibiting effects of zinc (Zn) and insulin that are co-present with hIAPP in islet cells, and the lipid membrane has been shown to significantly influence the aggregation kinetics. Increasing number of studies report the importance of developing small molecule inhibitors to suppress the hIAPP’s toxicity. Particularly, the ability of epigallocatechin-gallate (EGCG) to inhibit amyloid aggregation of a variety of amyloid peptide/proteins including hIAPP initiated numerous studies including the development of compounds to potentially treat amyloid diseases. In this study, by using a combination of thioflavin-T fluorescence and transmission electron microscopy experiments, we demonstrate a significant enhancement in EGCG’s efficiency, when mixed with Zn, to significantly suppress hIAPP amyloid aggregation both in presence and absence of lipid membrane. Circular dichroism experiments indicate the formation and stabilization of a helical structure of hIAPP in presence of EGCG:Zn complex. Our results also reveal the ability of EGCG or EGCG:Zn to suppress hIAPP’s cellular toxicity and that the ability of EGCG to chelate with Zn suppresses zinc’s cellular toxicity. We suggest that the reported results would be useful to develop strategies to trap hIAPP intermediates for further biophysical and structural studies, and also to devise approaches to abolish amyloid aggregation and cellular toxicity.

## 1 Introduction

Human-IAPP (hIAPP: human islet amyloid polypeptide, also known as amylin) is a 37 residue peptide hormone (Fig.S1A) that is co-secreted at a 1:100 molar ratio with insulin in response to blood glucose levels and normally contributes to satiety and the slowing of gastric emptying.[1] In type II diabetes, the increase in insulin production causes an increase in hIAPP amyloid deposits causing β cell damage. Biophysics studies have reported the aggregation of human-IAPP results in lipid membrane disruption following a two-step mechanism: fibril-independent pore formation and a fibril-dependent fragmentation of the lipid bilayer.[2] The presence of anionic lipids enhances monomeric hIAPP binding to the lipid membrane while the presence of curvature promotes aggregation.[3] Previous studies also reported that hIAPP forms toxic oligomers, which are alpha-helical.[4-6]

Because of the cellular toxicity that associates with amyloid formation, it is important to develop small molecule compounds that inhibit/prevent amyloid aggregation and toxicity.[7] One of the frequently investigated amyloid inhibitors is epigallocatechin-gallate (EGCG; the major catechin present in green tea) (Fig.S1B), a polyphenolic compound found in green tea extract.[8-12] EGCG has been shown to be an effective amyloid inhibitor for a variety of amyloid-forming peptides and proteins. In addition to small molecule modulation, metal ions were shown to play a role in amyloid aggregation. In pancreatic β cells Zn is among the highest in the body and Zn deficiency is a common symptom in type II diabetes.[13] Zinc has been shown to coordinate with His-18 of hIAPP and promotes the formation of off-pathway oligomers that are incompetent to grow to form amyloid fibers. Interestingly, Zn has a dual effect on hIAPP’s amyloid aggregation properties in which low concentrations of Zn inhibit fiber formation but high concentrations promote aggregation.[14-16] Here we studied the combined effect of EGCG and Zinc on alleviating the amyloid aggregation of hIAPP and its cellular toxicity. We report that Zn enhances the ability of EGCG to inhibit amyloid formation in the presence and absence of membranes. Because the membrane disruption induced by the amyloid aggregation of hIAPP is linked to islet cell death in type II diabetics, it is important to examine the efficiency of amyloid inhibition by small molecules in a membrane environment. By using a combination ThT based fluorescence and TEM, we show significant increase in the lag-time and reduction in the elongation rate, amorphous aggregate formation in presence of EGCG or EGCG:Zn complex. It is remarkable that EGCG:Zn complex stabilizes the helical structure of hIAPP. Our results also show that EGCG-Zn complex also increases the cell viability by suppressing zinc’s toxicity in cells.

## 2. Materials and Methods

### 2.1 Chemicals

Lyophilized hIAPP with amidated C-termini was purchased from Peptide Institute Inc (Osaka, Japan). In order to avoid undesired aggregation, hIAPP solution was carefully prepared according to the method published previously.[17] The concentration of hIAPP was spectroscopically determined based on a molar extinction coefficient of 1490 M^-1^ cm^-1^ at 280 nm. Two lipids in the powder form, 1-palmitoyl-2-oleoyl-sn-glycero-3-phosphocholine (POPC) and 1-palmitoyl-2-oleoyl-*sn*-glycero-3-phospho-rac-(1-glycerol) (POPG), were purchased from Avanti Polar Lipids, Inc. (Alabaster, AL). ThT was obtained from Wako Pure Chemical Industries, Ltd. ZnCl_2_, and EGCG and the other reagents were purchased from Nacalai Tesque (Kyoto, Japan).

### 2.2 Preparation of lipid vesicles

Large unilamellar vesicles (LUVs) with a diameter of 100 nm were prepared as described previously.[18] Briefly, POPC and POPG lipids were first dissolved in chloroform and were mixed at a 7:3 molar ratio for mimicking a mixed anionic/zwitterionic membrane system. The solvent was removed under a steam of nitrogen and dried overnight to generate a lipid film which was next rehydrated in 20 mM HEPES (pH 7.4) containing 50 mM NaCl. Lipid suspension was subjected to 5 rounds of freeze-thaw and the resulting solution was extruded 23 times through 100-nm polycarbonate membranes (Whatman, Clifton, NJ) using a mini-extruder (Avanti Polar Lipids, Inc., Alabaster, AL). Unless otherwise noted, LUVs which consisted of 250 μM lipids were used for the current study.

### 2.3 ThT assay for amyloid formation of hIAPP

Kinetics of amyloid fibrillation at 37 °C were monitored using 5 μM hIAPP in 20 mM HEPES buffer (pH 7.4) containing 5 μM ThT and 50 mM NaCl in the presence and absence of LUVs. Other effectors of hIAPP amyloid generation including Zn(II), EGCG, or EGCG:Zn(II) mixture with a 1:1 molar ratio were added to hIAPP sample solution at the desired concentration.

ThT-based monitoring was performed with a sealed 96-well microplate (Greiner-Bio-One, Tokyo, Japan). ThT fluorescence intensity was observed in a microplate reader (MTP-810, Corona Electric Co. Ibaraki, Japan) every 3 min following 10-sec shaking. The excitation and emission wavelengths were set at 450 and 490 nm, respectively. Kinetic parameters of amyloid formation were obtained by the following equation;[19]

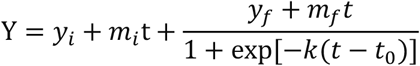

where *y*_i_ + *m*_i_*t* and *y*_f_ + *m*_f_*t* indicate initial and final baselines, respectively. *k* and *t* indicate the rate constant of elongation and time, respectively. *t*_0_ is the half-time when ThT fluorescence reaches 50% of its maximum intensity. The lag time was calculated with the relationship, lag time = *t*_0_ - 2(1/*k*).

### 2.4 Circular dichroism spectroscopy

Far-UV circular dichroism (CD) spectra were obtained at a concentration of 15 μM hIAPP dissolved in 20 mM HEPES containing 50 mM NaCl and 500 μM lipid in LUVs. Zn(II), EGCG, or EGCG:Zn(II) (1:1) complex was added to hIAPP sample solution at the desired concentration. CD measurements were performed on a Jasco J820 spectropolarimeter (Tokyo, Japan) using a quartz cuvette with a light path of 0.1 cm. A cell holder equipped with a water circulator was used to keep the sample temperature constant at 37 °C. Spectra were presented as mean residue ellipticity, [θ] (deg cm^2^ dmol^-1^), after subtraction of the solvent background.

### 2.5 Transmission electron microscopy

All transmission electron microscopy (TEM) images were obtained using a HITACHI H-7650 transmission microscope (Hitachi, Tokyo, Japan) at 20 °C with a voltage of 80 kV. Each sample solution was applied onto a collodion-coated copper grid (Nisshin EM Co., Tokyo, Japan), and negatively stained with 2% (weight/weight) ammonium molybdate as reported previously.[18]**2.6**

### 2.6 Preparation of EGCG:Zn(II) complex

EGCG:Zn(II) complex was prepared by mixing ZnCl_2_ and EGCG at the desired molar ratio in 20 mM HEPES buffer (pH 7.4) containing 50 mM NaCl. The absorption spectra of EGCG:Zn(II) complex were acquired using Hitachi U-3000 spectrometer (Hitachi, Tokyo, Japan).

### 2.7 Cell toxicity experiments

To prepare hIAPP fibrils for the cell proliferation assays, lyophilized hIAPP monomer was solubilized in Milli-Q water which was adjusted to pH 4 with HCl. Stock solutions were prepared by adding sample buffer (50 mM NaCl, 20 mM HEPES, pH 7.4) to a final concentration of 0.1 mg/ml. Peptide solutions at 100 μM were then treated with either 1 or 5 molar equivalents of EGCG:Zn(II). Peptide solutions at 100 μM were then treated with either 1 or 5 molar equivalents of EGCG:Zn(II). To form aggregates, peptides were incubated at room temperature with shaking at 750 rpm for 24 hours. RIN-5F cells (purchased from ATCC, cat# CRL-2058, 61465080) were grown in RPMI-1640 media with 2 mM L-glutamine supplemented with 10% fetal bovine serum in a humidified incubator at 37 °C with 5% CO_2_. Cells were kept between passage numbers 5-15. Cell were cultured in 10-cm cell culture dishes. For cell viability assays, 90μl of cells were dispensed to a total of 30,000 cells per well in 96 flat-bottom well trays and incubated for 24 hours at 37 °C with 5% CO_2_ prior the assay. Cell proliferation was measured by 3-(4,5-dimethylthiazol-2-yl)-2,5-di-phenyltetrazolium bromide (MTT) assay (CellTiter 96 Non-Radioactive Cell Proliferation Assay, Promega). 10 μL of peptides (freshly dissolved and aggregated) were added to each well to a final concentration of 1, 5 or 10 μM. After 24h incubation the cell proliferation was determined as described by the manufacturer. In short, 15 μL Dye solution was added to each well. After incubation at 37 °C for 3-3.5 hours. 100 μL of solubilization solution/stop mix was added. Plates were left at room temperature for 12 hours with gentle shaking. The absorbance was measured at 570 nm and 700 nm (background). Data was corrected for the background by subtracting the absorbance at 700nm from the absorbance at 570nm. The data was normalized with cells treated with 1% (w/v) SDS to 0% reduction, and cells treated with sample buffer to 100% reduction. Five technical replicates were used for each experiment. Values reported are the average of three independent experiments and the error is the standard deviation. Ordinary one-way ANOVA tests with Tukey’s multiple comparisons were performed in respect to non-treated cells (dotted line) and individual samples (solid line). ****p<0.0001, ***p*<*0.0002, and *p<0.05 indicate levels of significant differences.

### 2.8 Dynamic light scattering measurements

EGCG solutions were prepared in 20 mM HEPES buffer (pH 7.4) containing 50 mM NaCl and mixed well by gently vortex for 30 seconds. Solution mixture of EGCG and ZnCl_2_ were prepared at equimolar concentration and incubated for 15 minutes at 37 °C. The size distribution of EGCG/EGCG:Zn(II) complex solutions were measured by dynamic light scattering (DLS) at 37 °C. All DLS experiments were performed using a DynaPro NanoStar from Wyatt Technology (Santa Barbara, CA) with a 1-μL quartz cuvette and the average values over 20 independent scans were presented.

### 2.9 NMR experiments

One dimensional proton NMR spectra of hIAPP were recorded on a 600 MHz Bruker Avance III spectrometer equipped with a z-axis gradient cryogenic probe at 10 °C. The freshly dissolved hIAPP monomers in 20 mM HEPES (pH 6.5) containing 50 mM NaCl, 90% H_2_O, and 10% ^2^H_2_O were titrated with either 500 μM EGCG alone or 500 μM EGCG with 500 μM ZnCl_2_. The proton NMR spectra were processed using TopSpin 3.5 (Bruker BioSpin, Germany).

## 3. Results and Discussion

### 3.1 Effect of EGCG and Zn(II) on hIAPP aggregation in LUVs

The formation of EGCG:Zn(II) complex in solution (Fig.S2) and in 7:3 molar ratio of POPC:POPG LUVs (Fig.S3) was first characterized by UV-vis experiments. Experimental results reveal that EGCG is capable of binding to more than one unit of Zn(II) in solution and that the formation of EGCG:Zn(II) complex even in the membrane environment, which are in agreement with results reported in a previous study.[20] DLS experiments provided information on physical phase states of EGCG and EGCG:Zn(II) (Fig. S4). Hydrodynamic radius (*R*_H_) of EGCG was predominantly distributed at ∼7-8 nm while *R*_H_ of EGCG:Zn(II) displayed ∼5, ∼250, and ∼300-400 nm, which indicates that EGCG exists in an aggregated form while the EGCG:Zn(II) complex forms larger aggregates (Fig. S4).

Since both Zn(II) and EGCG are independently capable of interacting with hIAPP and suppress its fibril formation, the efficiencies of Zn(II), EGCG, and EGCG:Zn(II) complex to inhibit the amyloid aggregation of hIAPP in solution (Fig. S5) and in 7:3 POPC:POPG LUVs (Figures 1 and S6) were examined by ThT-based fluorescence experiments. The results reveal the remarkable ability of EGCG or EGCG:Zn(II) complex to inhibit the aggregation of hIAPP to form fibrils in solution and even in the presence of lipid bilayers. For a better comparison of the experimentally observed effects of Zn(II), EGCG, and EGCG:Zn(II) complex on the fibril formation of hIAPP, the observed changes in the lag time, elongation rate constant, and maximum ThT intensity are summarized in Fig. 1. Zn(II) alone slightly increases the lag time and slightly reduces the maximum ThT intensity without affecting the elongation rate constant (Fig.1A and D-F and Fig. S6). On the other hand, both EGCG and EGCG:Zn(II) mixture exhibited a considerable increase in the lag time as well as a reduction in both elongation rate constant and maximum ThT intensity (Fig.1B-F). For example, the elongation rate constant and ThT intensity are fully suppressed by 25-50 μM EGCG without an appreciable lag time (Fig. 1E). The complexation of EGCG with Zn(II) enhanced these effects in the presence of membrane (Fig. 1E) and also in solution (Fig. S4). In fact, 5 μM of 1:1 EGCG:Zn(II) mixture is sufficient to dramatically reduce both the elongation rate and ThT intensity in presence of LUVs (Fig.1E and F). At 2.5 μM of 1:1 EGCG:Zn(II) mixture, amyloid formation was completely prevented in solution as indicated by the remarkably low ThT fluorescence intensity (Fig. S5C), while a high ThT fluorescence intensity was still observed with a slow apparent elongation rate and prolonged lag time in the absence of Zn(II) (Fig. S5B).

**Figure 1.**
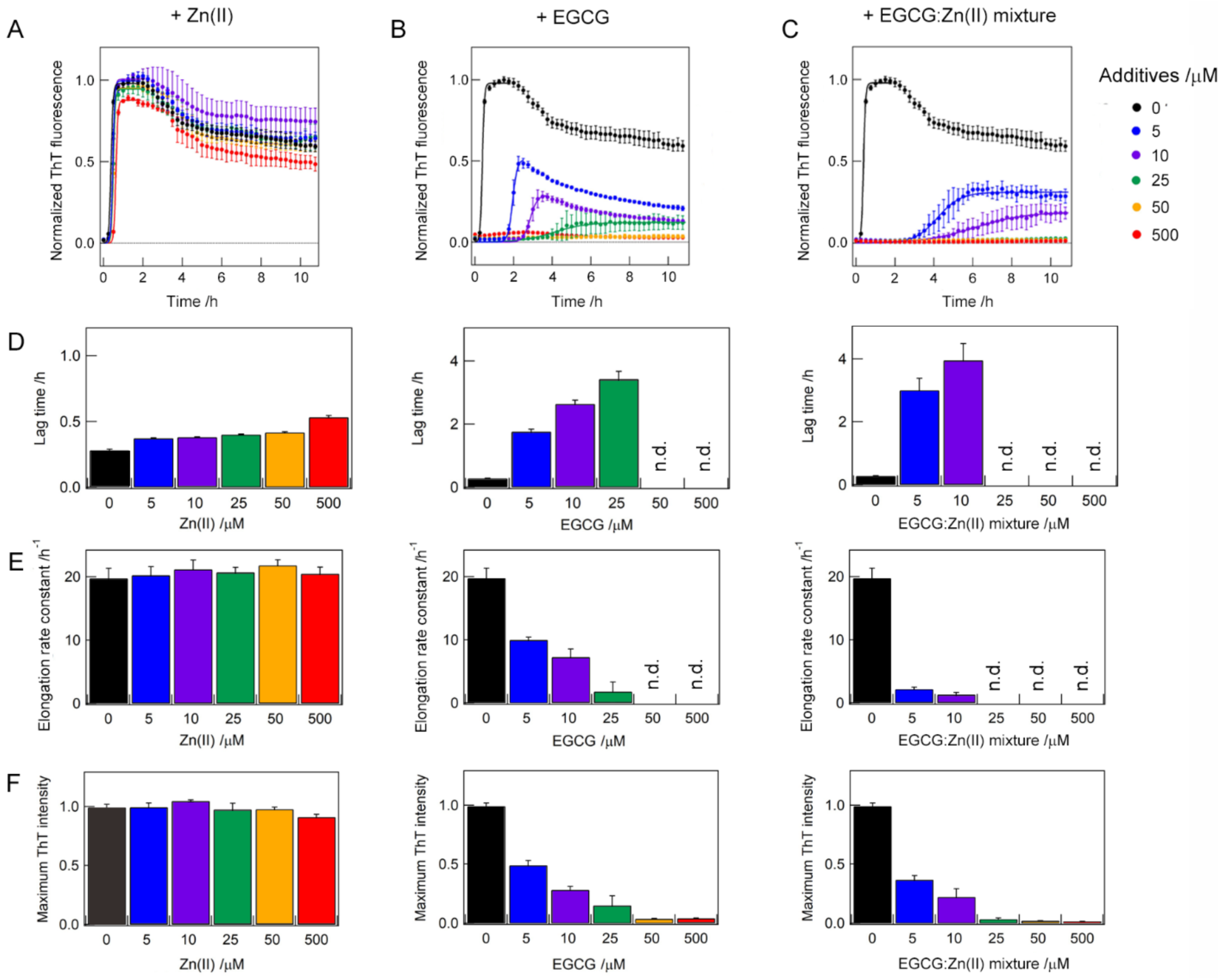
Inhibition of hIAPP amyloid formation in a lipid bilayer. (A-C) ThT fluorescence experiments showing the suppression of amyloid aggregation of hIAPP by Zn(II) (A), EGCG (B), EGCG:Zn(II) mixture (C) in the presence of 7:3 POPC:POPG LUVs. Lag time (D), elongation rate constant (E) and maximum ThT intensity (F) in the presence of Zn(II) (left), EGCG (middle) and EGCG:Zn(II) mixture (right). hIAPP of 5 μM was incubated at 37 °C and pH 7.4. Average values of three independent measurements are shown with error bars representing the standard deviation. Solid lines in A-D are the best-fit of experimental data points. “n.d.” denotes the concentration at which the best fit was not obtained due to very slow aggregation (i.e., very low ThT intensity).

Since EGCG is capable of interfering with the ThT binding to amyloid fibrils as reported elsewhere,[21,22] the ThT-based observation of the inhibition of hIAPP’s amyloid fibril formation is further confirmed by taking TEM images of the samples (Fig. 2). As shown in the TEM images, Zn(II) alone is capable of reducing the fibril density (Fig. 2A and B) in agreement with previous studies which showed that Zn(II) coordinates with His-18 and forms off-pathway complex that does not aggregate further to form amyloid fibrils.[15] In accordance with ThT results shown in Fig.1, Zn(II) alone cannot completely suppress the fibril formation of hIAPP. On the other hand, both EGCG and EGCG:Zn(II) are fairly efficient in fully suppressing the fibril formation by promoting the formation of amorphous aggregates of hIAPP (Fig. 2C-F). It is remarkable that the amount of EGCG required to completely suppress hIAPP fibril formation is dramatically reduced by the presence of Zn(II) as evidenced from experimental results shown in Figs. 1 (D-F) and S5 (B and C).

**Figure 2.**
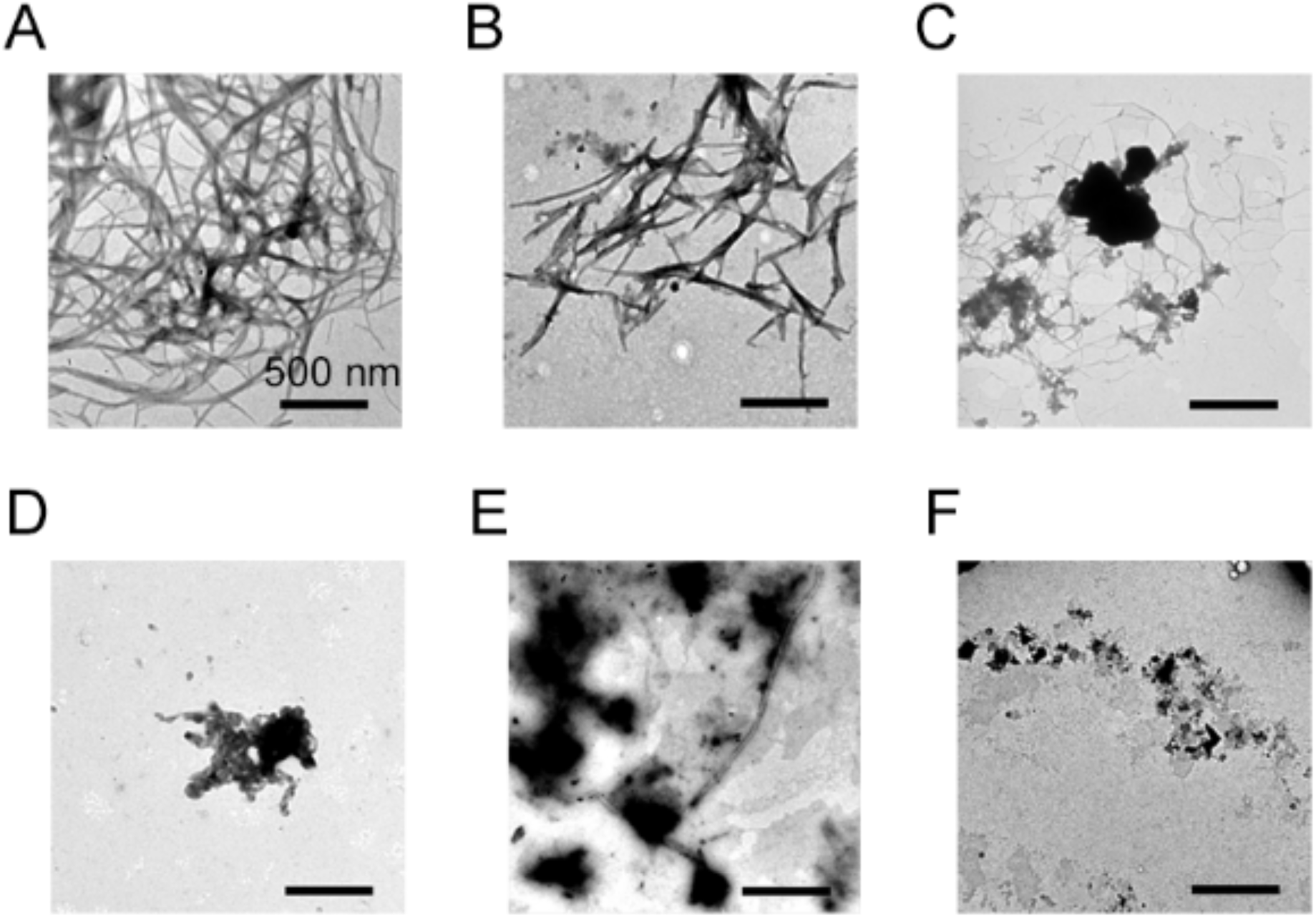
TEM images of hIAPP aggregates. Mature amyloid fibrils of hIAPP of 5 μM incubated for 12 hours with 7:3 POPC:POPG LUVs in the absence of additives (A) and in the presence of 500 μM Zn(II) (B), 5 μM EGCG (C), 500 μM EGCG (D), 5 μM EGCG:Zn(II) mixture (1:1 molar ratio) (E), and 500 μM EGCG:Zn(II) mixture (1:1 molar ratio) (F). The black scale bars represent 500 nm.

### 3.2 EGCG:Zn(II) complex reduces the β-sheet content of hIAPP aggregates

These ThT and TEM results were further confirmed by CD experiments (Fig. 3). hIAPP adopted initially a α-helical structure upon interacting with LUVs, characterized by two minima at 208 and 222 nm in the CD spectrum. Incubation of the solution generated typical CD spectra of the β-sheet structure which exhibits a single minimum ∼220 nm, revealing the conversion of helical structures to β-rich structures. Prediction of the content of the secondary structure using algorithm of BeStSel [23] indicated that α-helical structures of hIAPP almost transformed to β structured amyloid fibrils by showing ∼1-∼4% of helical structures and ∼30-∼40% of β structures, and that the presence of Zn(II) and/or EGCG did not affect markedly the content of the secondary structure of hIAPP amyloids (Table S1). The combination of CD and TEM results indicate that the formation of β-sheet structured amyloid fibrils of hIAPP in solution or in lipid bilayers both in the absence and presence of Zn(II), while the extent of fibril formation is reduced by the presence of Zn(II) (Figs. 2A,B and 3A,B). In addition, CD and TEM results indicate the formation of amorphous β-sheet aggregates in the presence of EGCG or EGCG:Zn(II) mixture as well as enhanced inhibiting capability of EGCG in the presence of Zn(II) (Figs. 2, 3, and S5D,E).

**Figure 3.**
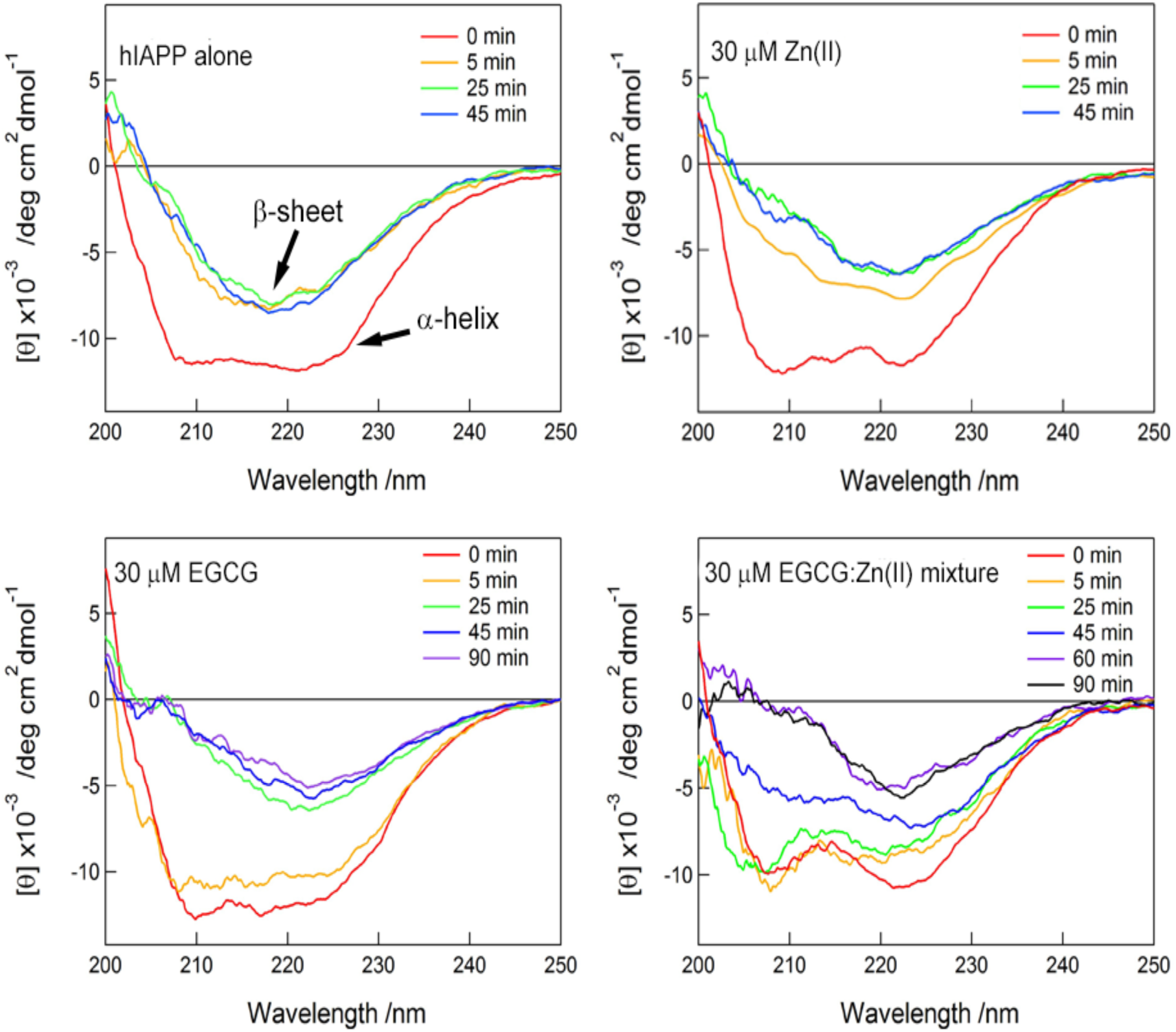
Real time CD spectra of hIAPP. Far-UV CD spectra of hIAPP at 15 μM in 7:3 POPC:POPG LUVs (A) and in the presence of 30 μM Zn(II) (B), 30 μM EGCG (C), and 30 μM of 1:1 molar ratio of EGCG:Zn(II) mixture (D). The formation and stabilization of a helical structure of hIAPP in presence of EGCG:Zn(II) complex for a longer duration is worth mentioning and could be important in enabling high-resolution structural studies of the intermediate.

### 3.3 Thermodynamic analysis of intermolecular interactions using isothermal titration calorimetry

As the inhibitory ability of EGCG in the absence and presence of Zn(II) inhibited hIAPP amyloid aggregation, we further examined intermolecular interactions using isothermal titration calorimetry (ITC) (Figs. 4 and S7). ITC thermograms showed exothermic binding reactions between hIAPP and either EGCG alone or EGCG:Zn(II) mixture (Fig. 4A and B), indicating intermolecular interactions. ITC analyses revealed that complex formation was both driven by the negative enthalpy change (Δ*H*) and the positive entropy change (Δ*S*), and reported the apparent binding stoichiometry ‘*n’* (the number of hIAPP binding to one EGCG or EGCG:Zn(II) complex). The values of ‘*n’* for hIAPP-EGCG and hIAPP-EGCG:Zn(II) binding systems were ∼0.5 and ∼0.8, respectively. The dissociation constant (*K*_d_) of hIAPP for EGCG (∼1.1 μM) is larger than that for EGCG:Zn(II) mixture (∼0.7 μM) to some extent. Thus, the results may implicate that EGCG:Zn(II) complex binds to hIAPP more tightly than EGCG and more hIAPP is capable of binding to EGCG:Zn(II) complex as compared to EGCG.

**Figure 4.**
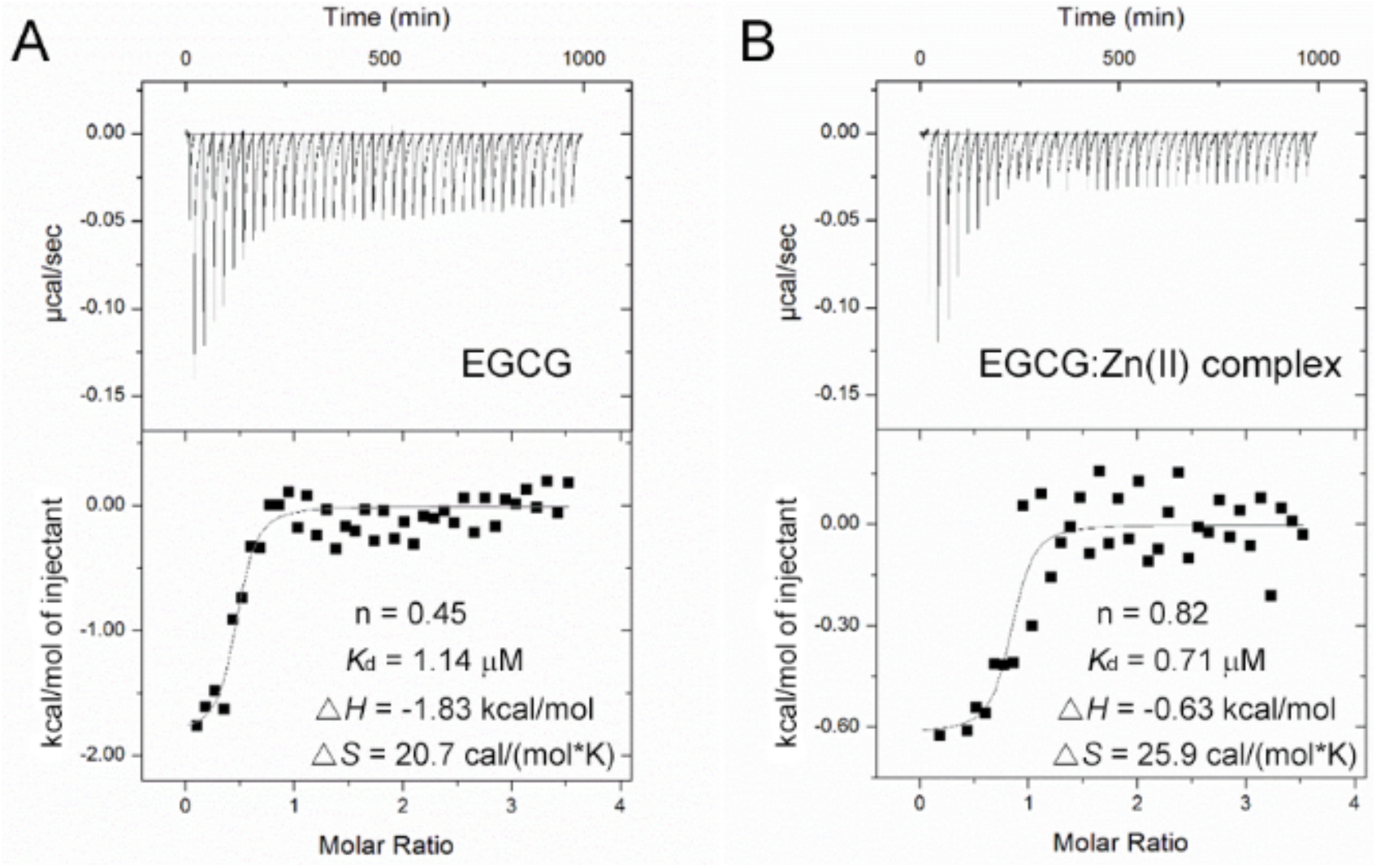
ITC measurements for the interactions between hIAPP and EGCG and EGCG:Zn(II) complex. (A and B) ITC thermograms (upper panel) and binding isotherms (lower panel) for the titration of 1 mM EGCG (A) or EGCG:Zn(II) mixture (B) into 60 μM hIAPP are shown.

### 3.4 One-dimensional proton NMR spectroscopy shows formation of large aggregates between hIAPP and EGCG:Zn(II) complex

A recent study reported morphologically distinct hIAPP aggregates in the presence of EGCG and suggested interactions between hIAPP and EGCG at the atomistic resolution.[12] Thus, to further characterize effects of Zn(II) on intermolecular interactions and phase states of EGCG as well as on hIAPP amyloid aggregation, we performed a series of one-dimensional proton NMR measurements (Fig. S7, left). NMR spectra showed line broadening of amide-proton resonances of hIAPP in the presence of EGCG (Fig.S7), which is in agreement with a recent NMR study.[12] As EGCG aggregates were detected by DLS measurements (Fig.S4) and photography (Fig.S7, right panel), binding of hIAPP (ITC analyses in Fig. 4) to EGCG aggregates was ascribed to line broadending. Interestingly, EGCG in the presence of the equimolar concentration of ZnCl_2_ significantly reduced the intensity of amide-proton NMR peaks (Fig.S7). In addition, hIAPP mixed with EGCG and ZnCl_2_ yielded in a precipitate whereas a turbid solution containing EGCG was observed (Fig.S7, right panel). Therefore, severe line broadening resulted from interactions between hIAPP and large aggregates of EGCG:Zn(II) complex as ITC analyses showed these molecular interactions.

Based on these results, we suggest that aggregate forms of EGCG serve as inhibitors of amyloid generation, and reason that Zn(II) binding reinforces intermolecular interactions among EGCG and thereby generating larger EGCG aggregates than EGCG aggregates without Zn(II) as observed in DLS measurements (Fig.S4). Indeed, other metal ions such as aluminum have also shown to enhance the antiamyloidogenic property of EGCG due to complex formation and aggregation.[12] Oligomers of 5,5,6,6-tetrachloro-1,1,3,3-tetraethylbenzimidazolylcarbocyanine iodide (JC-1), a fluorescent dye, bound to alpha-synuclein[24] and oligomers of polyphenolic compounds including EGCG and catechin exhibited higher potency to suppress amyloid fibrillation of amyloid-β and insulin than their monomeric forms.[25,26] Collectively, aggregated forms of EGCG:Zn(II) complex are expected to be excellent in inhibiting hIAPP amyloidogenesis in a more efficient way than EGCG by using better binding ability for hIAPP in terms of the binding affinity and the number of bound molecules as revealed in ITC analyses. In addition, it is worth noting that trapping helical structures of hIAPP by EGCG or EGCG:Zn(II) complex (Fig. 3) is also an important factor to suppress amyloidogenesis.

### 3.5 MTT assay

Because EGCG:Zn(II) complex is the most efficient in inhibiting amyloid formation of hIAPP, we tested whether it can also counteract the toxic effect of hIAPP by promoting cell viability. We used the reduction of MTT to evaluate the toxicity of hIAPP incubated with different molar ratios of EGCG:Zn(II) (Fig. 5). In agreement with previous studies,[5,27], monomeric hIAPP (1μM freshly dissolved) exhibit more toxicity than the fibrils of hIAPP, Fig 5A. hIAPP mixed with EGCG:Zn prior cell treatment (Fig.5C) or incubated for 24 hours (Fig 5B) showed increased cell proliferation in comparison to hIAPP monomers and fibrils alone. This increase in cell viability was statistically significant observed when comparing 10 μM hIAPP monomers or fibers alone with peptide treated with 1 or 5 equivalents of EGCG:Zn(II) (Table S2). However, when fresh peptide and EGCG:Zn(II) mixture were added to cells (Fig. 5B), the positive effect of the complex was less pronounced. Similar to treatment with EGCG only, treatment of cells with EGCG:Zn(II) caused an increase in cell proliferation (Figure S8 and S9). However, treatment with only Zn(II) caused a 40% reduction in cell viability (Figure S9). These results show the toxic nature of high concentrations of free Zn(II) and the ability of EGCG:Zn complex to negate it. Experiments were performed with hIAPP monomers and fibrils treated with either EGCG or Zn (Figure S10). A more pronounced increase in cell viability was seen with EGCG than with EGCG:Zn(II), and a more pronounced decrease was seen with Zn only. This shows the toxic impact that Zn alone can have on hIAPP toxicity and the redox active nature of EGCG. While the EGCG:Zn(II) complex is not as redox active as EGCG, it does show the importance of complex formation on rescuing Zinc toxicity.

**Figure 5.**
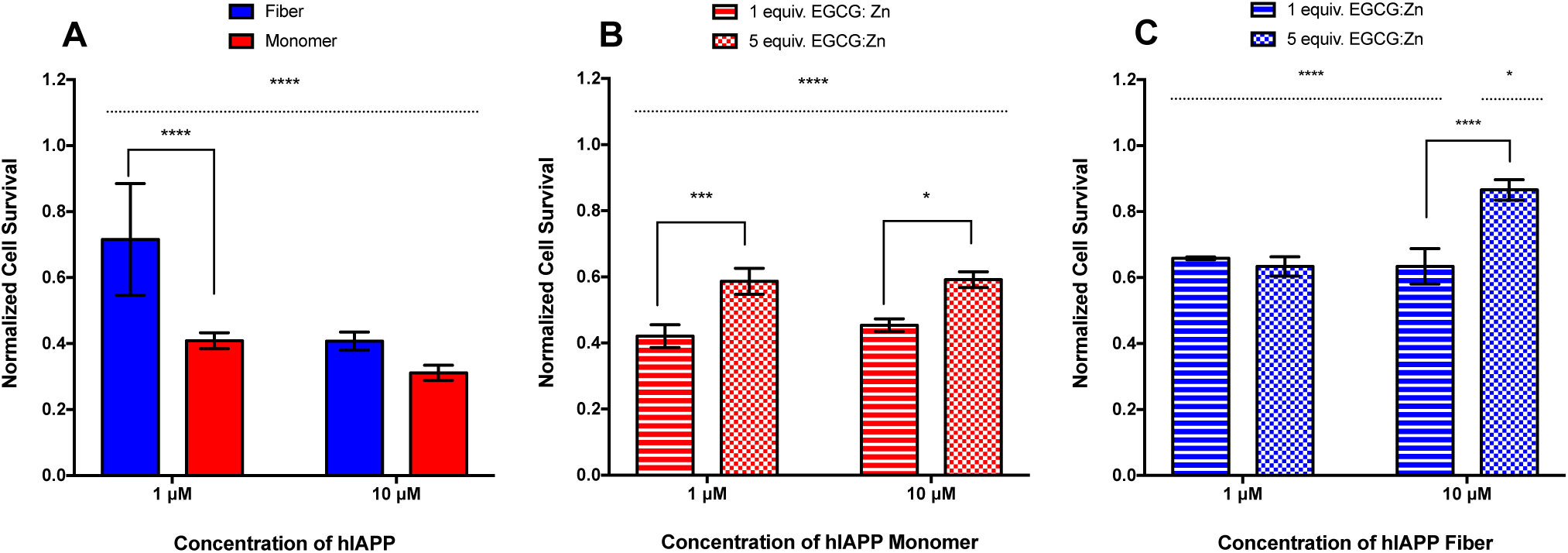
RIN-5F cell viability using MTT assay. (A) Cells treated with freshly prepared hIAPP monomer (red) and matured hIAPP fibrils (blue). (B) Freshly prepared hIAPP monomer added in conjunction with varying ratios of EGCG:Zn(II) mixture (red). (C) hIAPP fibrils grown for 24 hours with varying ratios of EGCG:Zn(II) mixture and then exposed to cells (blue). All samples were normalized using buffer and SDS as negative and positive controls, respectively. Ordinary one-way ANOVA tests with Tukey’s multiple comparisons were performed in respect to buffer treated cells (dotted line) and in respect to individual samples (solid line). ****p<0.0001, ***p*<*0.0002, and *p<0.05 indicate levels of significant differences. Data is shown as mean +/-SD with N=5 replicates.

Previous studies have reported the toxic nature of the EGCG:Zn(II) complex at higher concentrations on a different cell line than the one used in this study in absence any peptide [20,28]. Interestingly, Sun et. al. show that the cell permeability is enhanced by the presence of free Zn, inhibiting the cell growth. However, EGCG complexed with Zn(II) alleviates the toxic effect of the metal. While zinc can induce off-pathway toxic amyloid aggregates formation [29], the EGCG:Zn(II) mixture was found to protect cells in our system (Figure S8). Taken together, the deficiency of Zn(II) in β-cell granules involving in the T2D development [30] could be supplemented in the form of EGCG:Zn(II) complex. This can increase the cell proliferation as well as deliver Zn to deficient cells via increased membrane permeability, also can inhibit hIAPP aggregation that is linked to β-cell failure, giving this complex a unique dual effect.

## 4. Conclusions

EGCG has been shown to be an efficient inhibitor of amyloid aggregation by a variety of amyloidogenic peptides and proteins, and has also been shown to remodel amyloid fibers.[31-42] Although many small molecules can inhibit amyloid aggregation in solution, it is important to evaluate the efficiency of such compounds in the presence of membrane which could be directly related to their ability to suppress cell toxicity.[17,43,44] In this study, we have successfully demonstrated the ability of EGCG to suppress the amyloid fibril formation of hIAPP both in solution and in presence of membrane by using a combination of ThT fluorescence, TEM and CD experiments. In addition, the reported results demonstrate that the presence of Zn(II) reduced the amount of EGCG required to completely inhibit hIAPP’s fibril formation. These results are further confirmed by the cell toxicity measurements that show the ability of EGCG:Zn(II) complex to significantly suppress hIAPP’s toxicity. In fact, EGCG suppress zinc’s toxicity to cells as observed from the significant decrease in cell viability in the presence of EGCG:Zn complex. Our findings provide insightful information for the future development of therapeutic small molecules targeting amyloid diseases.

While the EGCG:Zn complex has been shown to be efficient in suppressing hIAPP’s amyloid fibril formation, it would be useful to investigate the structural effects of zinc on EGCG and the factors that enhances the efficacy of EGCG:Zn complex. Our NMR experiments showed that the presence of zinc enhanced the aggregation of EGCG-hIAPP to result in the precipitation of the sample. A systematic analysis of the aggregated and/or precipitated samples by magic angle spinning solid-state NMR experiments would provide insights into Zn-EGCG and EGCG:Zn-hIAPP interactions. It is remarkable that EGCG:Zn complex stabilizes helical oligomeric hIAPP intermediates within the elongated lag-time as revealed by CD experiments. High-resolution structures of such intermediates can be determined by NMR experiments, although the intermediate structures may have to be stabilized by cooling (or freezing) the sample for MAS solid-state NMR experiments. Structures of these intermediates would be useful to better understand the aggregation pathways as well as the mechanism of amyloid inhibition by EGCG or EGCG:Zn.

## Acknowledgements

This study was supported by NIH (AG048934 to A.R.) and a Grant-in-Aid for Young Scientists (B) (15K18518 and 25870407) (to Y.-H.L.).

## References

1. Westermark P, Andersson A, Westermark GT: Islet amyloid polypeptide, islet amyloid, and diabetes mellitus. Physiol Rev 2011, 91:795–826.

2. Brender JR, Salamekh S, Ramamoorthy A: Membrane disruption and early events in the aggregation of the diabetes related peptide IAPP from a molecular perspective. Acc Chem Res 2012, 45:454–462.

3. Sciacca MF, Brender JR, Lee DK, Ramamoorthy A: Phosphatidylethanolamine enhances amyloid fiber-dependent membrane fragmentation. Biochemistry 2012, 51:7676–7684.

4. Nanga RP, Brender JR, Vivekanandan S, Ramamoorthy A: Structure and membrane orientation of IAPP in its natively amidated form at physiological pH in a membrane environment. Biochim Biophys Acta 2011, 1808:2337–2342.

5. Abedini A, Plesner A, Cao P, Ridgway Z, Zhang J, Tu LH, Middleton CT, Chao B, Sartori DJ, Meng F, et al.: Time-resolved studies define the nature of toxic IAPP intermediates, providing insight for anti-amyloidosis therapeutics. Elife 2016, 5.

6. Birol M, Kumar S, Rhoades E, Miranker AD: Conformational switching within dynamic oligomers underpins toxic gain-of-function by diabetes-associated amyloid. Nat Commun 2018, 9:1312.

7. Doig AJ, Derreumaux P: Inhibition of protein aggregation and amyloid formation by small molecules. Curr Opin Struct Biol 2015, 30:50–56.

8. Popovych N, Brender JR, Soong R, Vivekanandan S, Hartman K, Basrur V, Macdonald PM, Ramamoorthy A: Site specific interaction of the polyphenol EGCG with the SEVI amyloid precursor peptide PAP(248-286). J Phys Chem B 2012, 116:3650–3658.

9. Hora M, Carballo-Pacheco M, Weber B, Morris VK, Wittkopf A, Buchner J, Strodel B, Reif B: Epigallocatechin-3-gallate preferentially induces aggregation of amyloidogenic immunoglobulin light chains. Sci Rep 2017, 7:41515.

10. Lorenzen N, Nielsen SB, Yoshimura Y, Vad BS, Andersen CB, Betzer C, Kaspersen JD, Christiansen G, Pedersen JS, Jensen PH, et al.: How epigallocatechin gallate can inhibit alpha-synuclein oligomer toxicity in vitro. J Biol Chem 2014, 289:21299–21310.

11. Andrich K, Bieschke J: The Effect of (-)-Epigallo-catechin-(3)-gallate on Amyloidogenic Proteins Suggests a Common Mechanism. Adv Exp Med Biol 2015, 863:139–161.

12. Franko A, Rodriguez Camargo DC, Boddrich A, Garg D, Rodriguez Camargo A, Rathkolb B, Janik D, Aichler M, Feuchtinger A, Neff F, et al.: Epigallocatechin gallate (EGCG) reduces the intensity of pancreatic amyloid fibrils in human islet amyloid polypeptide (hIAPP) transgenic mice. Sci Rep 2018, 8:1116.

13. Foster MC, Leapman RD, Li MX, Atwater I: Elemental composition of secretory granules in pancreatic islets of Langerhans. Biophys J 1993, 64:525–532.

14. Salamekh S, Brender JR, Hyung SJ, Nanga RP, Vivekanandan S, Ruotolo BT, Ramamoorthy A: A two-site mechanism for the inhibition of IAPP amyloidogenesis by zinc. J Mol Biol 2011, 410:294– 306.

15. Brender JR, Hartman K, Nanga RP, Popovych N, de la Salud Bea R, Vivekanandan S, Marsh EN, Ramamoorthy A: Role of zinc in human islet amyloid polypeptide aggregation. J Am Chem Soc 2010, 132:8973–8983.

16. Brender JR, Krishnamoorthy J, Messina GM, Deb A, Vivekanandan S, La Rosa C, Penner-Hahn JE, Ramamoorthy A: Zinc stabilization of prefibrillar oligomers of human islet amyloid polypeptide. Chem Commun (Camb) 2013, 49:3339–3341.

17. Pithadia AS, Bhunia A, Sribalan R, Padmini V, Fierke CA, Ramamoorthy A: Influence of a curcumin derivative on hIAPP aggregation in the absence and presence of lipid membranes. Chem Commun (Camb) 2016, 52:942–945.

18. Terakawa MS, Yagi H, Adachi M, Lee YH, Goto Y: Small liposomes accelerate the fibrillation of amyloid beta (1-40). J Biol Chem 2015, 290:815–826.

19. Nielsen L, Khurana R, Coats A, Frokjaer S, Brange J, Vyas S, Uversky VN, Fink AL: Effect of environmental factors on the kinetics of insulin fibril formation: elucidation of the molecular mechanism. Biochemistry 2001, 40:6036–6046.

20. Sun SL, He GQ, Yu HN, Yang JG, Borthakur D, Zhang LC, Shen SR, Das UN: Free Zn(2+) enhances inhibitory effects of EGCG on the growth of PC-3 cells. Mol Nutr Food Res 2008, 52:465–471.

21. Meng F, Abedini A, Plesner A, Verchere CB, Raleigh DP: The flavanol (-)-epigallocatechin 3-gallate inhibits amyloid formation by islet amyloid polypeptide, disaggregates amyloid fibrils, and protects cultured cells against IAPP-induced toxicity. Biochemistry 2010, 49:8127–8133.

22. Suzuki Y, Brender JR, Hartman K, Ramamoorthy A, Marsh EN: Alternative pathways of human islet amyloid polypeptide aggregation distinguished by (19)f nuclear magnetic resonance-detected kinetics of monomer consumption. Biochemistry 2012, 51:8154–8162.

23. Micsonai A, Wien F, Kernya L, Lee YH, Goto Y, Refregiers M, Kardos J: Accurate secondary structure prediction and fold recognition for circular dichroism spectroscopy. Proc Natl Acad Sci U S A 2015, 112:E3095–3103.

24. Lee JH, Lee IH, Choe YJ, Kang S, Kim HY, Gai WP, Hahn JS, Paik SR: Real-time analysis of amyloid fibril formation of alpha-synuclein using a fibrillation-state-specific fluorescent probe of JC-1. Biochem J 2009, 418:311–323.

25. Nie RZ, Zhu W, Peng JM, Ge ZZ, Li CM: A-type dimeric epigallocatechin-3-gallate (EGCG) is a more potent inhibitor against the formation of insulin amyloid fibril than EGCG monomer. Biochimie 2016, 125:204–212.

26. Hayden EY, Yamin G, Beroukhim S, Chen B, Kibalchenko M, Jiang L, Ho L, Wang J, Pasinetti GM, Teplow DB: Inhibiting amyloid β-protein assembly: Size-activity relationships among grape seed-derived polyphenols. J Neurochem 2015, 135:416–430.

27. Sciacca MF, Kotler SA, Brender JR, Chen J, Lee DK, Ramamoorthy A: Two-step mechanism of membrane disruption by Abeta through membrane fragmentation and pore formation. Biophys J 2012, 103:702–710.

28. Yang J, Yu H, Sun S, Zhang L, Das UN, Ruan H, He G, Shen S: Mechanism of free Zn(2+) enhancing inhibitory effects of EGCG on the growth of PC-3 cells: interactions with mitochondria. Biol Trace Elem Res 2009, 131:298–310.

29. Lee MC, Yu WC, Shih YH, Chen CY, Guo ZH, Huang SJ, Chan JCC, Chen YR: Zinc ion rapidly induces toxic, off-pathway amyloid-β oligomers distinct from amyloid-β derived diffusible ligands in Alzheimer’s disease. Sci Rep 2018, 8:4772.

30. Jayawardena R, Ranasinghe P, Galappatthy P, Malkanthi R, Constantine G, Katulanda P: Effects of zinc supplementation on diabetes mellitus: a systematic review and meta-analysis. Diabetol Metab Syndr 2012, 4:13.

31. Palhano FL, Lee J, Grimster NP, Kelly JW: Toward the molecular mechanism(s) by which EGCG treatment remodels mature amyloid fibrils. J Am Chem Soc 2013, 135:7503–7510.

32. Lopez del Amo JM, Fink U, Dasari M, Grelle G, Wanker EE, Bieschke J, Reif B: Structural properties of EGCG-induced, nontoxic Alzheimer’s disease Aβ oligomers. J Mol Biol 2012, 421:517–524.

33. Hyung SJ, DeToma AS, Brender JR, Lee S, Vivekanandan S, Kochi A, Choi JS, Ramamoorthy A, Ruotolo BT, Lim MH: Insights into antiamyloidogenic properties of the green tea extract (-)- epigallocatechin-3-gallate toward metal-associated amyloid-β species. Proc Natl Acad Sci U S A 2013, 110:3743–3748.

34. Huang R, Vivekanandan S, Brender JR, Abe Y, Naito A, Ramamoorthy A: NMR characterization of monomeric and oligomeric conformations of human calcitonin and its interaction with EGCG. J Mol Biol 2012, 416:108–120.

35. Fusco G, Sanz-Hernandez M, Ruggeri FS, Vendruscolo M, Dobson CM, De Simone A: Molecular determinants of the interaction of EGCG with ordered and disordered proteins. Biopolymers 2018:e23117.

36. Jha NN, Kumar R, Panigrahi R, Navalkar A, Ghosh D, Sahay S, Mondal M, Kumar A, Maji SK: Comparison of α-Synuclein Fibril Inhibition by Four Different Amyloid Inhibitors. ACS Chem Neurosci 2017, 8:2722–2733.

37. Ahmed R, Melacini G: A solution NMR toolset to probe the molecular mechanisms of amyloid inhibitors. Chem Commun (Camb) 2018, 54:4644–4652.

38. Tu LH, Young LM, Wong AG, Ashcroft AE, Radford SE, Raleigh DP: Mutational analysis of the ability of resveratrol to inhibit amyloid formation by islet amyloid polypeptide: critical evaluation of the importance of aromatic-inhibitor and histidine-inhibitor interactions. Biochemistry 2015, 54:666–676.

39. Xu ZX, Ma GL, Zhang Q, Chen CH, He YM, Xu LH, Zhou GR, Li ZH, Yang HJ, Zhou P: Inhibitory Mechanism of Epigallocatechin Gallate on Fibrillation and Aggregation of Amidated Human Islet Amyloid Polypeptide. Chemphyschem 2017, 18:1611–1619.

40. Mo Y, Lei J, Sun Y, Zhang Q, Wei G: Conformational Ensemble of hIAPP Dimer: Insight into the Molecular Mechanism by which a Green Tea Extract inhibits hIAPP Aggregation. Sci Rep 2016, 6:33076.

41. Wang J, Yamamoto T, Bai J, Cox SJ, Korshavn KJ, Monette M, Ramamoorthy A: Real-time monitoring of the aggregation of Alzheimer’s amyloid-β via. Chem Commun (Camb) 2018, 54:2000–2003.

42. Pithadia A, Brender JR, Fierke CA, Ramamoorthy A: Inhibition of IAPP Aggregation and Toxicity by Natural Products and Derivatives. J Diabetes Res 2016, 2016:2046327.

43. Malishev R, Shaham-Niv S, Nandi S, Kolusheva S, Gazit E, Jelinek R: Bacoside-A, an Indian Traditional-Medicine Substance, Inhibits β-Amyloid Cytotoxicity, Fibrillation, and Membrane Interactions. ACS Chem Neurosci 2017, 8:884–891.

44. Engel MF, vandenAkker CC, Schleeger M, Velikov KP, Koenderink GH, Bonn M: The polyphenol EGCG inhibits amyloid formation less efficiently at phospholipid interfaces than in bulk solution. J Am Chem Soc 2012, 134:14781–14788.

